# Adjuvant conditioning shapes the adaptive immune response and promotes trained immunotolerance via NLRP3/IL-1

**DOI:** 10.1101/2024.09.06.611736

**Authors:** Thais Boccia, Weikang Pan, Victor Fattori, Rodrigo Cervantes-Díaz, Michael S. Rogers, Ivan Zanoni, Alex G. Cuenca

## Abstract

Trained immunity enhances responsiveness of the innate immune system upon restimulation. Although adjuvants are used to enhance immune responses, we showed that repeated administration of alum, termed adjuvant conditioning (AC), establishes an immunosuppressive environment that delays allogeneic graft rejection by expanding myeloid-derived suppressor cells (MDSCs). Here, we show that AC-induced MDSCs suppress antigen specific adaptive responses both in vitro and in vivo, and that the immunosuppression is abolished in the absence of NLRP3 and IL-1 signaling. Allogeneic pancreatic islets transplanted into AC-treated NLRP3^-/-^ mice are not protected, demonstrating that AC requires NLRP3 signaling. Finally, AC also has an immunosuppressive effect on human cells. Overall, our data show that AC establishes an immunosuppressive milieu via the NLRP3/IL-1 axis, leading to trained immunosuppression, or trained tolerance. Our findings give a potent mandate to explore the possibility to target the NLRP3/IL-1 pathway as a new promising strategy to condition transplant recipients and promote allograft tolerance.

## Introduction

The utilization of adjuvants in the context of immunization potentiates the adaptive immune response driven by T and B cells. Aluminum salts (hereafter alum) are among the major adjuvants utilized in human vaccine formulations to potentiate the immune response against an antigen. Nevertheless, ‘non-specific’ antigen-independent immunization with alum before immunization has been associated with immunosuppression of the adaptive immune response, with lower total and specific IgM and IgG production (Schmoeckel et al., 2018). Similarly to alum, systemic administration of CpG-rich DNA oligonucleotides prior to immunization has been linked to reduced T cell expansion and diminished ovalbumin-specific cytotoxic T lymphocyte (CTL) activity (Wingender et al., 2006). Treatment of mice with an exopolysaccharide from Bacillus subtilis demonstrated alleviation of graft-versus-host disease severity (Kalinina et al., 2021), highlighting the role of Toll-like receptor (TLR) activation in establishing an immunosuppressive milieu in the context of allotransplant. We previously demonstrated that, when alum is employed as an adjuvant for non-specific immunization, a strategy termed adjuvant conditioning (AC), it induced the expansion of myeloid-derived suppressor cells (MDSCs) (Ge et al., 2023). AC-induced MDSCs suppressed T cell proliferation in vitro and delayed rejection of allogeneic pancreatic islets. These findings underscore the potential of adjuvant utilization as a pre-treatment strategy to enhance immune suppression preceding transplantation.

MDSCs represent a subset of immature myeloid cells originating from the bone marrow during systemic inflammation or cancer progression (Serafini et al., 2006; Melani et al., 2003; Bronte et al., 2016). These cells are comprised of various populations such as monocytes and granulocytes MDSCs and can be detected in the peripheral blood and spleens of both mice and humans (Almand et al., 2001; Young and Lathers, 1999; Ge et al., 2023). Monocyte-MDSCs (M-MDSCs) are CD11b+Ly6C^high^ cells, whereas polymorphonuclear (PMN)-MDSCs exhibit CD11b+Ly6C^low^Ly6G+ markers (Damuzzo et al., 2015; Dolcetti et al., 2008). MDSCs play a significant role in fostering cancer angiogenesis and tumor growth (Ahn and Brown, 2008; Yang et al., 2008), contributing to anti-tumor drug resistance and metastasis (Yang et al., 2008; Shojaei et al., 2007; Marvel and Gabrilovich, 2015; Veglia et al., 2021). Moreover, MDSCs have shown promise in therapeutic applications within transplantation models in mice (Ge et al., 2023; Dugast et al., 2008; Yang et al., 2016; He et al., 2015, 2016; Zhang et al., 2018). Among the suppressive mechanisms employed by MDSCs are the release of nitric oxide and prostaglandin E2, potent inhibitors of T cell responses (Rodriguez et al., 2005; Donkor et al., 2009; Mao et al., 2013; Navasardyan and Bonavida, 2021; Sreeramkumar et al., 2012). Additionally, there is a growing recognition of the correlation between inflammasome-dependent IL-1 and the expansion, mobilization, and suppressive function of tumor-induced MDSCs (Bunt et al., 2006, 2007; Papafragkos et al., 2022; Tengesdal et al., 2021; Elkabets et al., 2010; Luo et al., 2022; Shi et al., 2022; Koehn et al., 2019). This association can be attributed to the dual role of this pro-inflammatory cytokine in promoting resistance to infection while also contributing to inflammation control.

The term trained immunity describes an enhanced responsiveness of the innate immune system upon restimulation, resembling the immunological memory traditionally associated only with adaptive immune cells (Bonilla and Oettgen, 2010; Netea et al., 2020). Microbial ligand training of the innate system can confer non-specific protection against subsequent challenges in mice (Luzio and Williams, 1978; Krahenbuhl et al., 1981; Muñoz et al., 2010; Marakalala et al., 2013; Ribes et al., 2014), suggesting adaptive characteristics within the innate system. So far innate immune training has been explored only in the context of enhanced responses following exposure to microbial products. So far, there are no evidence that innate training can lead to immunosuppression. Here, we explored the possibility to utilize alum to drive a suppressive immune training, that we termed trained immunosuppression or trained tolerance, extending allograft survival.

## Results

### Adjuvant conditioning (AC) shapes the adaptive immune response in vivo

AC expands MDSCs, which inhibit T cell proliferation in vitro, as well as prolong the survival of pancreatic islet transplantation in an allogeneic setting (Ge et al., 2023). To assess the impact of adjuvant conditioning on the development of the adaptive immune response in vivo, we AC-treated, or not, mice and subsequently immunized the mice with ovalbumin (OVA) after OT-II cells adoptive transfer (Fig 1A). The number of OVA-specific CD4 T cells was increased in immunized-only mice compared to mice that received AC and immunization (Fig S1, Fig 1B. Additionally, Th1-skewed CD4+Vα+Tbet+ cells as well as Th2-skewed CD4+Vα+GATA3+ cells were significantly decreased in AC-immunized, compared to immunized-only, mice (Fig 1C-D).

**Figure 1:**
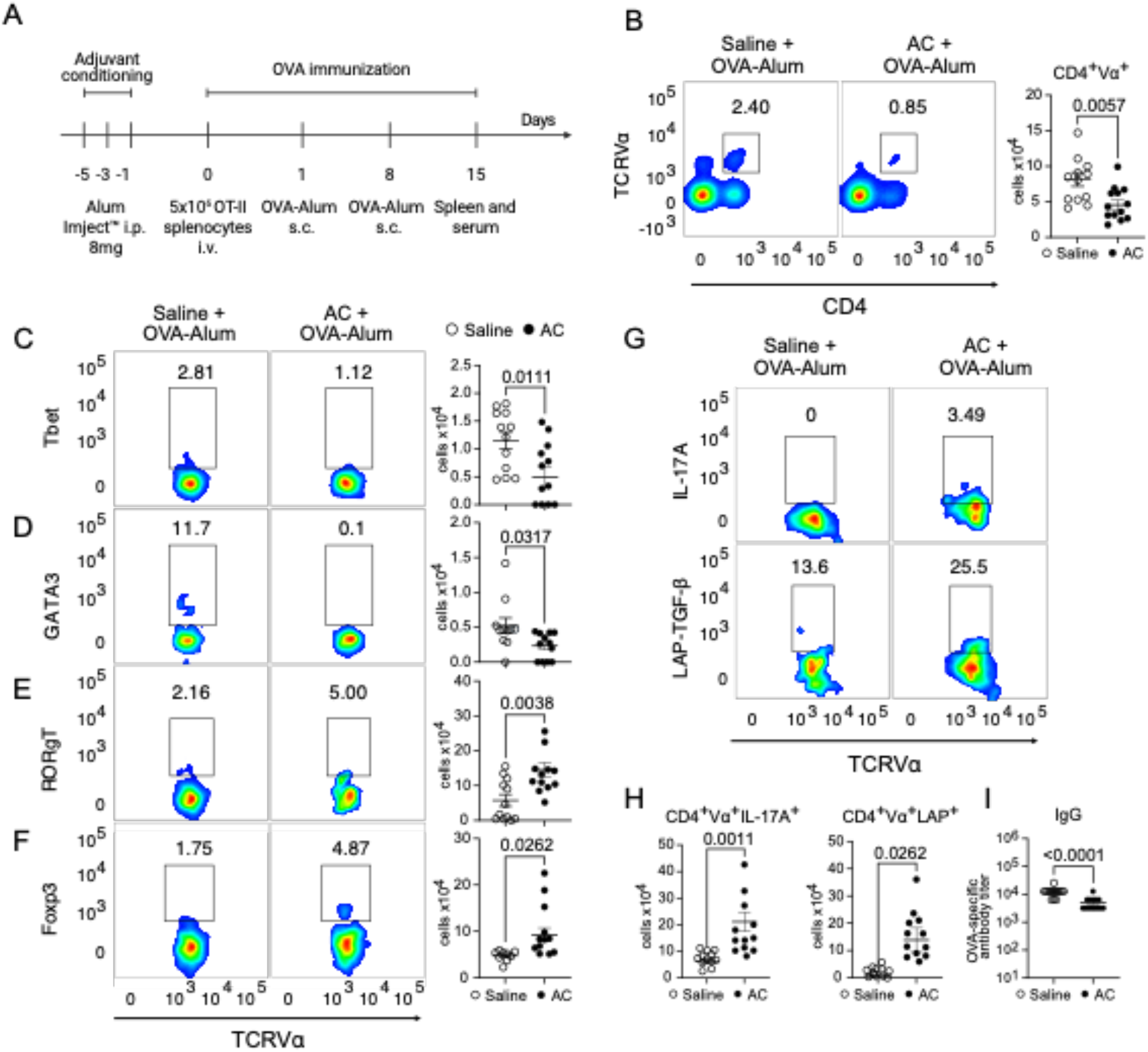
Adjuvant conditioning (AC) shapes the adaptive immune response *in vivo*. **A:** Experimental design. 8-10-week-old male or female C57BL/6 mice were injected intraperitoneally (i.p.) with either Alum Imject™ (8mg in 200µL) or saline three times, every other day. 24 hours after the last injection, mice were adoptively transferred with 5×10^6^ OT-II splenocytes intravenously, and 24 hours after the transfer, mice were immunized subcutaneously with 10 μg OVA adsorbed in 1.5mg of Alum Imject™, in a homologous prime (day 1) and boost (day 8) protocol. Seven days after the boost (day 15), mice were euthanized, and spleens were analyzed for OVA-specific Th1, Th2, Th17, and Treg total numbers. **B:** Flow cytometry plots and total number of OVA-specific T cells, evidenced by the expression of CD4^+^Vα^+^ in splenocytes from mice treated with either saline or alum before the immunization. Flow cytometry plots and total number of OVA-specific Th1 **(C)**, Th2 **(D)**, Th17 **(E)**, and Tregs **(F)**, evidenced by the expression of CD4^+^Vα^+^ Tbet^+^, CD4^+^Vα^+^GATA3^+^, CD4^+^Vα^+^RORγT^+^, and CD4^+^Vα^+^Foxp3^+^ in splenocytes from mice treated with either saline or alum before the immunization. **G:** Flow cytometry plots and total number **(H)** of OVA-specific Th17 and Tregs, evidenced by the expression of CD4^+^Vα^+^IL-17A and CD4^+^Vα^+^LAP^+^ in splenocytes treated with either saline or alum before the immunization and further cultured with OVAp (2μg/mL) *in vitro* for 48h in the presence of brefeldin-A. **I:** OVA-specific IgG titer in serum from mice treated with either saline or alum before the immunization. Data shown represent three or more experiments and are expressed as mean±SEM; Student t-test was used for analysis; P values are indicated in each graph. All statistical analyses were performed using GraphPad Prism™.

This was also accompanied by decreased production of OVA-specific IgG (Fig 1I).

Of note, AC-immunized mice exhibited enhanced differentiation of Th17 cells and regulatory T cells (Tregs) (CD4+Vα+RORγT+ and CD4+Vα+Foxp3+ respectively), compared to immunized-only (Fig 1E-F). This differentiation pattern was further confirmed when splenocytes were restimulated in vitro with OVA, as AC-immunized mice displayed elevated production of interleukin-17A (IL-17A) and transforming growth factor-beta (TGF-β) (CD4+Vα+IL-17A+ and CD4+Vα+LAP+) (Fig 1G-H). These results demonstrate that AC treatment dampens the adaptive immune response.

### NLRP3 activation is involved in the effects of AC on the adaptive immune response in vivo

Alum is a widely employed adjuvant in immunization protocols (Zhao et al., 2023), and its activity has been shown to depend on NLRP3 activation (Hornung et al., 2008; Eisenbarth et al., 2008). Indeed, we confirmed that NLRP3-deficient mice exhibited reduced expansion of OVA-specific CD4 T cells and lower levels of OVA-specific IgG compared to C57BL/6 mice when immunized with OVA in the presence of alum (Fig S2B-C), as previously shown (Li et al., 2008; Eisenbarth et al., 2008).

To assess whether AC effects involve NLRP3 activation, we performed AC with alum and immunized mice with OVA in the presence of resiquimod (R848) as an adjuvant (Tomai et al., 2007) (Fig 2A). We found that AC dampened CD4 T cell responses to OVA, as well as OVA-specific IgG and IgE antibody levels, also in the presence of resiquimod (Fig 2B-D and Fig S2C). The behavior of AC-treated mice was lost, or significantly decreased in mice that lack NLRP3 (Fig 2B-D and Fig S2C). We also observed that AC decreased Th1 and Th2 responses under these experimental conditions (Fig S2D and Fig 2E). Furthermore, AC followed by immunization of mice deficient in NLRP3 heightened differentiation not only of Th1 and Th2 cells but also of Th17 cells and Tregs (Fig S2D and Fig 2E). Overall, these data demonstrate a pivotal role for the NLRP3 inflammasome in driving the immunosuppression that follows AC.

**Figure 2:**
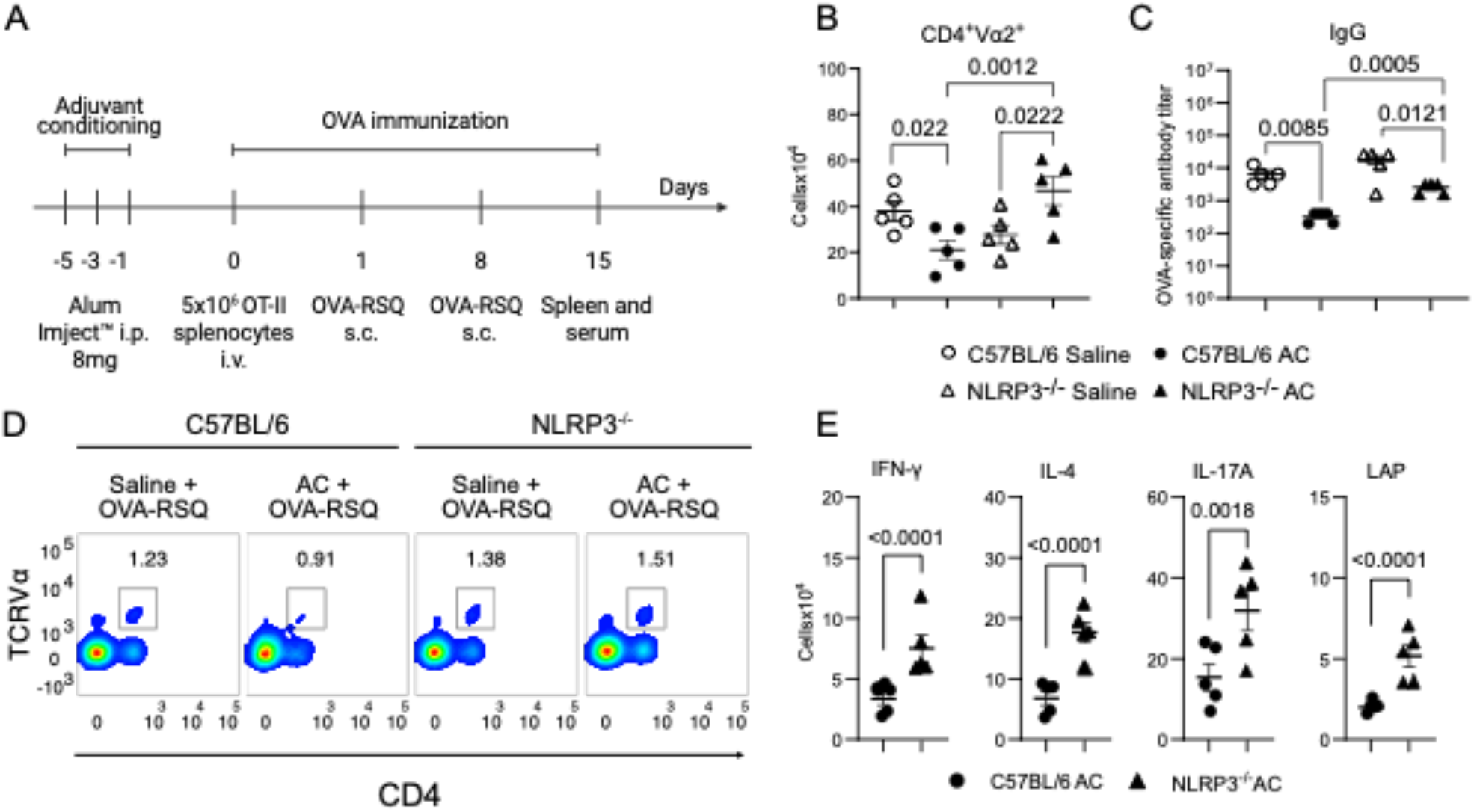
NLRP3 activation required for the effects of adjuvant conditioning in the adaptive immune response. **A**. Experimental design. 8-10-week-old male or female C57BL/6 or NLRP3^-/-^ mice were injected intraperitoneally (i.p.) with either Alum Imject™ (8mg in 200µL) or saline three times, every other day. 24 hours after the last injection, mice were adoptively transferred with 5×10^6^ OT-II splenocytes intravenously, and 24 hours after the transfer, mice were immunized with 10 μg OVA adsorbed in Resiquimod (R848) (OVA-RSQ) (50µg), in a homologous prime (day 1) and boost (day 8) protocol. Seven days after the boost (day 15), mice were euthanized, and spleens were analyzed for OVA-specific Th1, Th2, Th17, and Treg total numbers. **B:** Total number of OVA-specific T cells, evidenced by the expression of CD4^+^Vα^+^ in splenocytes from mice treated with either saline or alum before the immunization. **C:** OVA-specific IgG titer in serum from mice treated with either saline or alum before the immunization. **D:** Flow cytometry plots of OVA-specific T cells, evidenced by the expression of CD4^+^Vα^+^ in splenocytes from mice treated with either saline or alum before the immunization. **E:** Total number of OVA-specific Th1, Th2, Th17 and Tregs, evidenced by the expression of CD4^+^Vα^+^IFN-γ^+^, CD4^+^Vα^+^IL-4^+^, CD4^+^Vα^+^IL-17A^+^ and CD4^+^Vα^+^LAP^+^ respectively in splenocytes treated with either saline or alum before the immunization and further cultured with OVAp (2μg/mL) *in vitro* for 48h in the presence of brefeldin-A. Data shown represent three or more experiments, and are expressed as mean±SEM; Student t-test was used for analysis; P values are indicated in the graphs. All statistical analyses were performed using GraphPad Prism™.

### NLRP3-dependent IL-1 signaling is involved in the effects of AC on the adaptive immune response in vivo

Although extensive literature exists regarding the inflammatory effects of IL-1, recent studies have unveiled its capacity to foster the expansion of immunosuppressive cells such as MDSCs and Tregs, consequently promoting tumor growth (Arpaia et al., 2015; Papafragkos et al., 2022; Popovic et al., 2017; Elkabets et al., 2010). Notably, we found that NLRP3 deficiency impedes the production of both IL-1α and IL-1 β following AC (Fig S2A). To determine whether the AC-driven immunosuppression involves IL-1 signaling, we used Anakinra to block IL-1 signaling in the context of AC, followed by adoptive transfer of OT-II splenocytes and alum-OVA immunization. Our findings indicate that IL-1 blockade restored the proliferation of OVA-specific CD4 T cells (Fig 3A-B), facilitated Th1 and Th2 differentiation (Fig S2E and Fig 3D), and restored OVA-specific IgG and IgE production (Fig 2C and Fig S2F). Intriguingly, the elevation in Th17 and Treg differentiation persisted (Fig S2E and Figure 3D), suggesting that the Th17/Treg phenotype induced by the conditioning is independent of IL-1 signaling.

**Figure 3:**
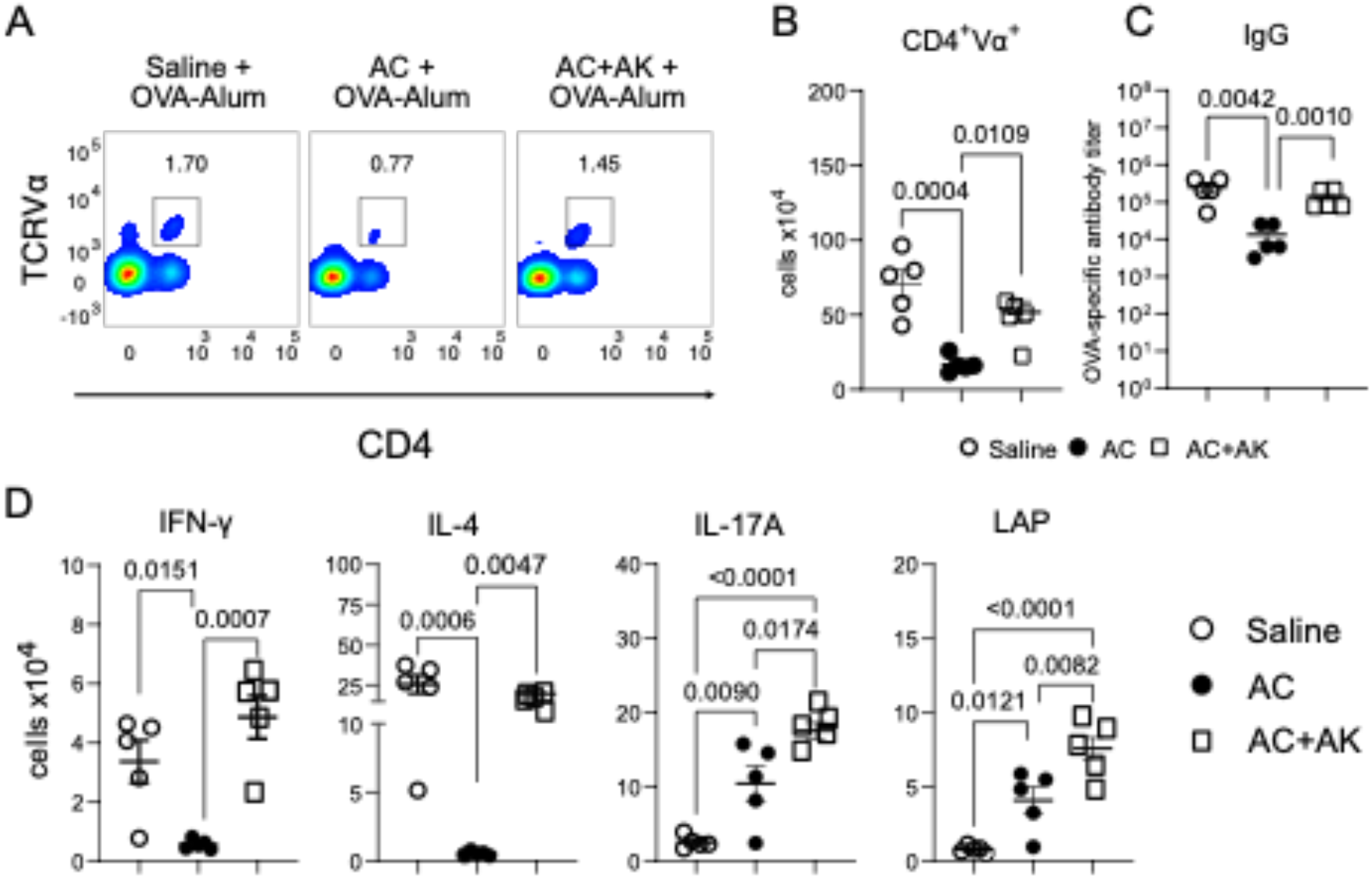
NLRP3-dependent IL-1 signaling is involved in the effects of adjuvant conditioning in the adaptive immune response. **A:** Flow cytometry plots and total number **(B)** of OVA-specific T cells, evidenced by the expression of CD4^+^Vα^+^ in splenocytes from mice treated with either saline, alum or alum ^+^ Anakinra (30mg/kg) (AC^+^AK) before the immunization. **C:** OVA-specific IgG titer in serum from mice treated with either saline, alum or alum + Anakinra (30mg/kg) (AC+AK) before the immunization. **D:** Total number of OVA-specific Th1, Th2, Th17 and Tregs, evidenced by the expression of CD4^+^Vα^+^IFN_-γ_^+^, CD4^+^Vα^+^IL-4^+^, CD4^+^Vα^+^IL-17A^+^ and CD4^+^Vα^+^LAP^+^ in splenocytes treated with either saline, alum or alum + Anakinra (30mg/kg) (AC+AK) before the immunization and further cultured with OVAp (2μg/mL) *in vitro* for 48h in the presence of brefeldin-A. Data shown represent three or more experiments, and are expressed as mean±SEM; Student t-test was used for analysis; P values are indicated in the graphs. All statistical analyses were performed using GraphPad Prism™.

### Adjuvant conditioning induces the reprogramming of myeloid cells to a trained immunosuppression phenotype

In agreement with our previous findings that showed the capacity of AC to expand MDSCs (Ge et al., 2023), AC led to an increase in total splenocytes and spleen weight (Fig S3A-B), and to the expansion of CD11b+Ly6C+ and CD11b+Ly6G+ cells (Fig S3C-D, and Fig 4A). To test the immunosuppressive activity of MDSCs differentiated upon AC treatment, we isolated splenic CD11b+GR1+ cells from saline or AC-treated mice were isolated and treated with LPS (200ng/mL) in vitro. CD11b+GR1+ cells from conditioned mice produced reduced levels of TNF-α and IL-6 compared to the cells isolated from saline-treated mice (Fig 4B-C). Additionally, these cells showed elevated levels of IL-10 and nitric oxide (Fig 1D-E) compared to their saline-treated counterparts, indicating an immature neutrophil/monocyte suppressor programming (Papafragkos et al., 2022; Kalafati et al., 2020). In functional assays, AC-induced splenic CD11b+GR1+ cells demonstrated potent suppression of CD4 T cell proliferation, with efficacy extending up to a 1:4 MDSC:T cell ratio (Fig 4F-G) and induced increased CD4 T cell death when compared to saline MDSCs at 1:1 ratio (Fig 4H).

**Figure 4:**
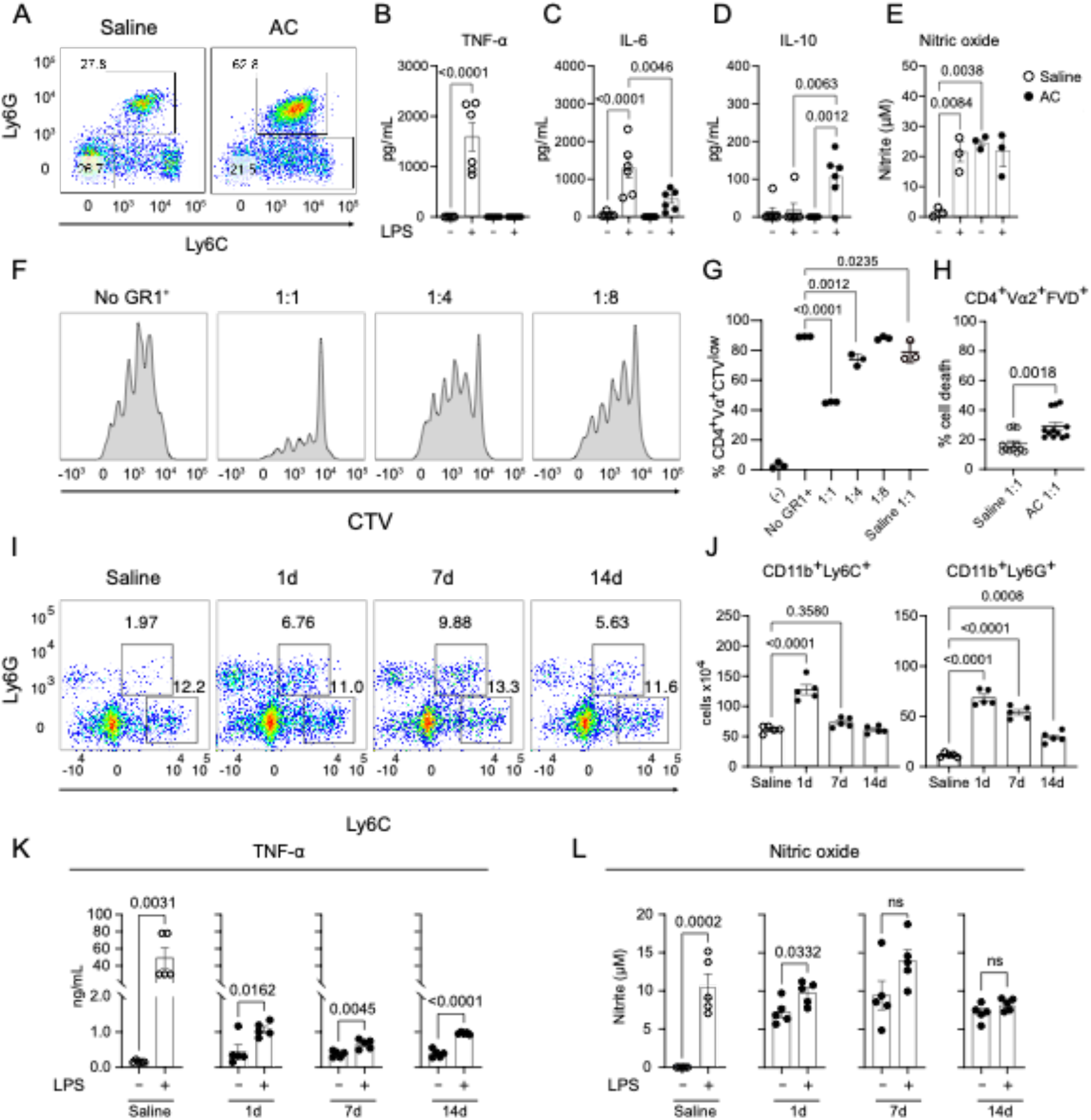
Adjuvant conditioning induces the reprogramming of myeloid cells to a trained immunosuppression phenotype. **A:** Flow cytometry plots of CD11b^+^Ly6C^+^ and CD11b^+^Ly6G^+^ cells in the spleen 24h after saline or alum injections. **B:** TNF-α, IL-6 **(C)**, IL-10 **(D)**, and nitric oxide **(E)** production from CD11b^+^GR1^+^ cells isolated from saline or AC-treated mice cultured with either medium or LPS (200ng/mL) for 24h. **F:** Histograms of CD4^+^Vα^+^ cells in a suppression assay using isolated CD11b^+^GR1^+^cells. Isolated MDSCs from spleens from AC-treated mice were cultured with OVAp (1µg/mL), C57BL/6 splenocytes (1×10^5^) and isolated OT-II naïve CD4 T cells (2×10^5^) in the ratios (MDSC : T cell) indicated in the graph. **G:** Percentage of CD4^+^Vα^+^CTV^low^ cells in a suppression assay using isolated CD11b^+^GR1^+^ cells and OT-II CD4 T cells. **H:** Percentage of CD4^+^Vα^+^FVD^+^ cells in a suppression assay using isolated CD11b^+^GR1^+^ cells and OT-II CD4 T cells. **I:** Flow cytometry plots of CD11b^+^Ly6C^+^ and CD11b^+^Ly6G^+^ cells in the spleen 1 day, 7 days, and 14 days after saline or alum injections. **J:** Total count of CD11b^+^Ly6C^+^ and CD11b^+^Ly6G^+^ cells in the spleen 1 day, 7 days, and 14 days after saline or alum injections. **K:** TNF-α, and nitric oxide **(L)** production from CD11b^+^GR1^+^ cells isolated 1 day, 7 days, and 14 days after saline or alum injections cultured with either medium or LPS (200ng/mL) for 24h. Data shown represent three or more experiments and are expressed as mean±SEM; Student t-test was used for analysis; P values are indicated in each graph. All statistical analyses were performed using GraphPad Prism™.

Furthermore, the suppressive effect persisted up to 14 days following conditioning. This is evidenced by sustained levels of CD11b+Ly6C+ and CD11b+Ly6G+ cells (Fig 4I-J), with a gradual increase in PDL-1, and sustained IDO expression (Fig S3F-H). When isolated, CD11b+GR1+ cells exhibit low TNF-αα production upon LPS administration (Fig 4K), while nitric oxide production was present regardless of LPS stimulation (Fig 4L). These results demonstrate that AC leads to a long-lasting reprogramming of CD11b+GR1+ cells that acquire a trained immunosuppressive phenotype.

### AC-induced MDSC expansion and suppressor function require NLRP3 activation and IL-1 signaling

Since there are reports of NLRP3 and IL-1 involvement in MDSC expansion and suppressor function in tumor models (Papafragkos et al., 2022), we next assessed the role of the NLRP3/IL-1 axis on AC-induced MDSC expansion and suppressor function. AC-treated NLRP3-/-mice revealed a notable decrease in the expansion of CD11b+Ly6C+ and CD11b+Ly6G+ populations compared to their wild-type counterpart (Fig 5A). Additionally, we observed a compromised suppressor function of AC-induced CD11b+GR1+ cells in the absence of NLRP3 (Fig 5B and Fig S3J), which was also observed for the ability to induce T CD4 cell death (Fig S3K).

**Figure 5:**
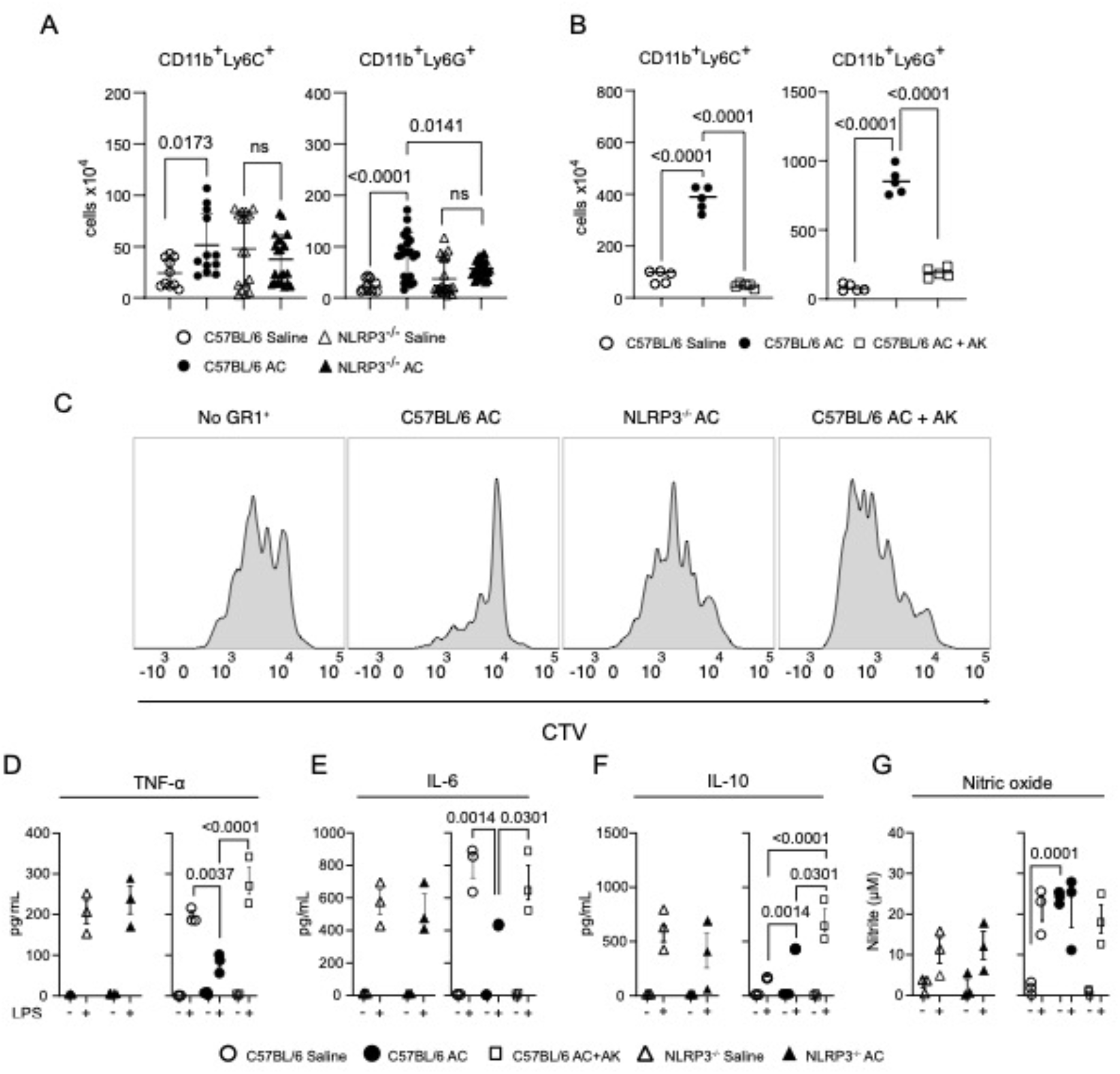
AC-induced MDSCs expansion and suppressor function requires NLRP3 activation and IL-1 signaling. **A:** Total number of CD11b^+^Ly6C^+^ and CD11b^+^Ly6G cells in the spleen 24h after saline or alum injections in C57BL/6 and NLRP3 deficient mice. **B:** Total number of CD11b^+^Ly6C^+^ and CD11b^+^Ly6G cells in the spleen 24h after saline, alum, or alum ^+^ Anakinra (30mg/kg) (AC+AK) injections. **C:** Histograms of CD4^+^Vα^+^ cells in a suppression assay using isolated CD11b^+^GR1^+^ from C57BL/6 and NLRP3 deficient mice treated with saline, alum, or alum + Anakinra (30mg/kg) (AC+AK). **D:** TNF-α, IL-6 **(E)**, IL-10 **(F)**, and nitric oxide **(G)** production from CD11b^+^GR1^+^ cells isolated from C57BL/6 or NLRP3^-/-^ mice injected either with saline, alum, or alum ^+^ Anakinra (30mg/kg) (AC+AK), cultured with either medium or LPS (200ng/mL) for 24h. Data shown represent three or more experiments, and are expressed as mean±SEM; Student t-test was used for analysis; P values are indicated in each graph. All statistical analyses were performed using GraphPad Prism™.

To further explore the requirement for IL-1 signaling downstream of NLRP3 activation for MDSC expansion and suppressor function, we administered Anakinra during the conditioning protocol. Remarkably, a similar pattern to the one observed with NLRP3 deficiency was observed. the expansion of both CD11b+Ly6C+ and CD11b+Ly6G+ populations was reduced, accompanied by impaired suppressor abilities when compared to AC-induced MDSCs derived from wild-type mice (Fig 5C-D and Fig S3L-M). This indicates that NLRP3 activation and subsequent IL-1 signaling during AC are crucial for the expansion and suppressor function of MDSCs, as previously shown for tumor-induced MDSCs (Papafragkos et al., 2022). Furthermore, we found no evidence of trained immunosuppression in AC-treated NLRP3-/-CD11b+GR1+ cells, or in the presence of IL-1 signaling blockade. Instead, we observed increased production of TNF-αα and IL-6 production, alongside diminished nitric oxide release in NLRP3-/-CD11b+GR1+ cells treated with LPS (Fig 5D-G), similarly saline controls, reinforcing the role of NLRP3 activation for the reprogramming of CD11b+GR1+ cells to a trained immunosuppressive phenotype.

### AC-induced MDSCs inhibit the adaptive immune response in vivo via NLRP3 activation and IL-1 signaling

To investigate the impact of AC-induced MDSCs on shaping the adaptive immune response in vivo, we adoptively transferred CD11b+GR1+ cells from AC or saline-treated mice into C57BL/6 recipients and immunized the mice (Fig 6A). Mice that received AC-induced MDSCs exhibited impaired expansion of OVA-specific CD4 T cells compared to those receiving CD11b+GR1+ cells from saline-injected mice (Fig 6B-C). Moreover, recipients of AC-induced MDSCs showed diminished levels of OVA-specific IgG production (Fig 6D), demonstrating that MDSCs induced during AC exposure also suppress the adaptive immune response in vivo. To explore the involvement of the NLRP3/IL-1 axis in MDSCs suppressor function in vivo, we adoptively transferred AC or saline-induced CD11b+GR1+ cells isolated from NLRP3-deficient mice, or C57BL/6 mice treated with Anakinra during the AC-treatment. Mice receiving NLRP3-deficient AC or saline-induced MDSCs demonstrated comparable expansion of OVA-specific CD4 T cells in both percentage and absolute numbers to control immunized mice, and similar results were obtained in mice receiving CD11b+GR1+ cells from Anakinra-treated donors (Fig 6E-F). Furthermore, OVA-specific IgG production was restored in mice that received NLRP3-deficient AC or saline-induced CD11b+GR1+ cells, but not those receiving cells from Anakinra-treated donors (Fig 6G), suggesting that IgG production is sensitive to the activity of NLRP3, but not to IL-1 signaling. Overall, these data underscore the essential role of the NLRP3/IL-1 axis in the suppressive function of AC-induced MDSCs in vivo.

**Figure 6:**
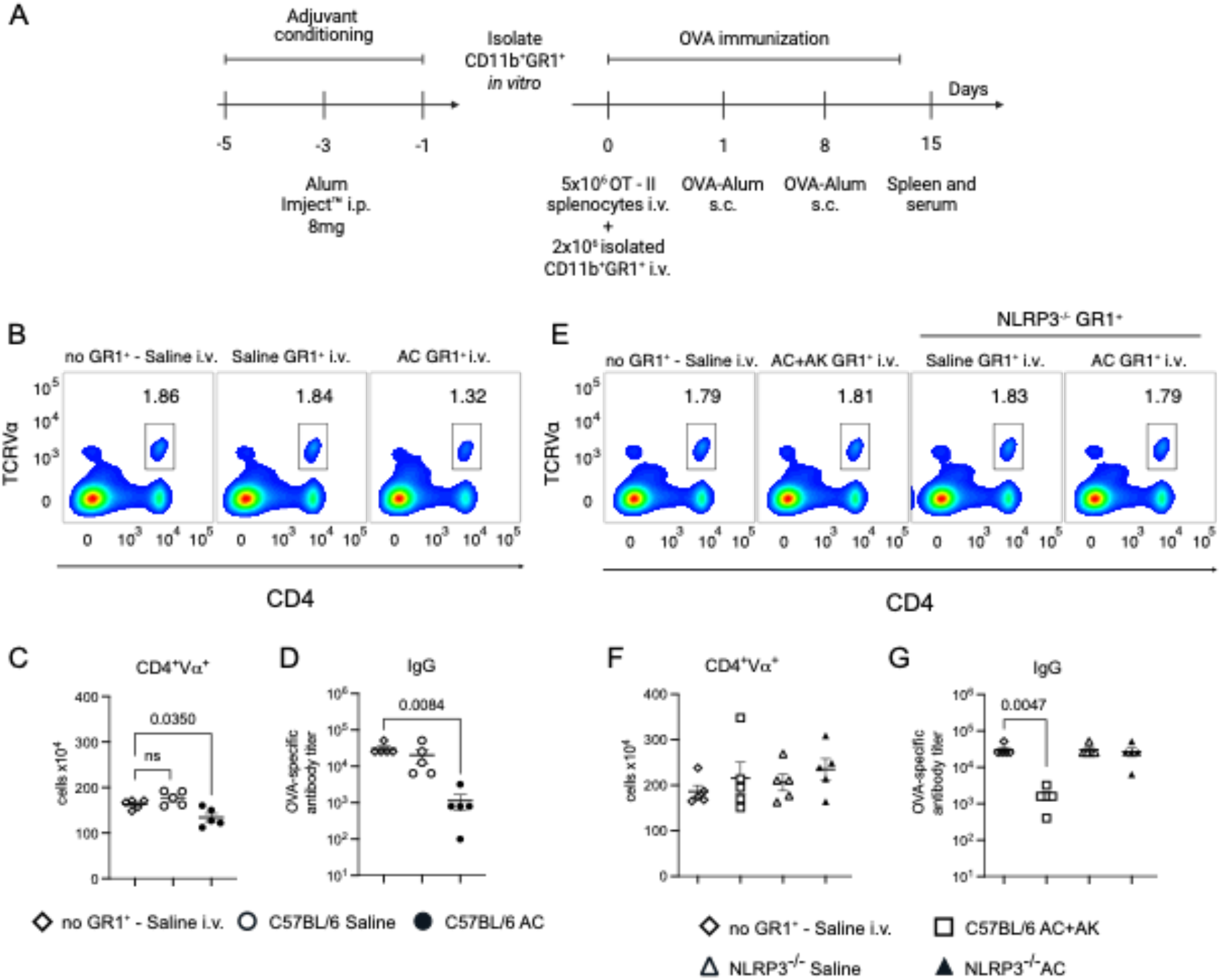
AC-induced MDSCs inhibition of the adaptive immune response *in vivo* requires NLRP3 activation and IL-1 signaling. **A:** Experimental design. Briefly, 2×10^6^ CD11b^+^GR1^+^ cells isolated from either AC, alum + Anakinra (30mg/kg) (AC+AK) or saline-injected C57BL/6 or NLRP3^-/-^ mice were adoptively transferred to C57BL/6 mice before the immunization protocol. **B:** Flow cytometry plots and total number of OVA-specific T cells **(C)**, evidenced by the expression of CD4^+^Vα^+^ in splenocytes from mice adoptively transferred with CD11b^+^GR1^+^ cells isolated from either AC, or saline-injected C57BL/6 before the immunization protocol. **D:** OVA-specific IgG titer in serum from mice adoptively transferred with CD11b^+^GR1^+^ cells isolated from either AC, or saline-injected C57BL/6 before the immunization protocol. **E:** Flow cytometry plots and total number of OVA-specific T cells **(F)**, evidenced by the expression of CD4^+^Vα^+^ in splenocytes from mice adoptively transferred with CD11b^+^GR1^+^ cells isolated from either alum + Anakinra (30mg/kg) (AC+AK) or saline-injected C57BL/6 or NLRP3^-/-^ mice before the immunization protocol. **G:** OVA-specific IgG titer in serum from mice adoptively transferred with CD11b^+^GR1^+^ cells isolated from either alum + Anakinra (30mg/kg) (AC^+^AK) or saline-injected C57BL/6 or NLRP3^-/-^ mice before the immunization protocol. Data shown represent three or more experiments and are expressed as mean±SEM; Student t-test was used for analysis; P values are indicated in the graphs. All statistical analyses were performed using GraphPad Prism™.

### AC-induced in vivo allogeneic tolerance requires NLRP3 activation

We previously demonstrated that adjuvant conditioning prolongs the survival of allogeneic pancreatic islet transplants through the expansion of MDSCs (Ge et al., 2023). Since we found that AC-induced MDSCs expansion and suppression function are dependent on NLRP3, we explored whether this effect extends to allogeneic responses in vivo and performed allogeneic islet transplantation in C57BL/6 and NLRP3^-/-^ mice that were AC-treated, or not (Fig 7A). Our findings revealed that AC significantly delays allogeneic islet rejection in wild-type but not in NLRP3-deficient mice (Fig 7B), underscoring the crucial role of NLRP3 in promoting the observed protective effect of AC in vivo.

**Figure 7:**
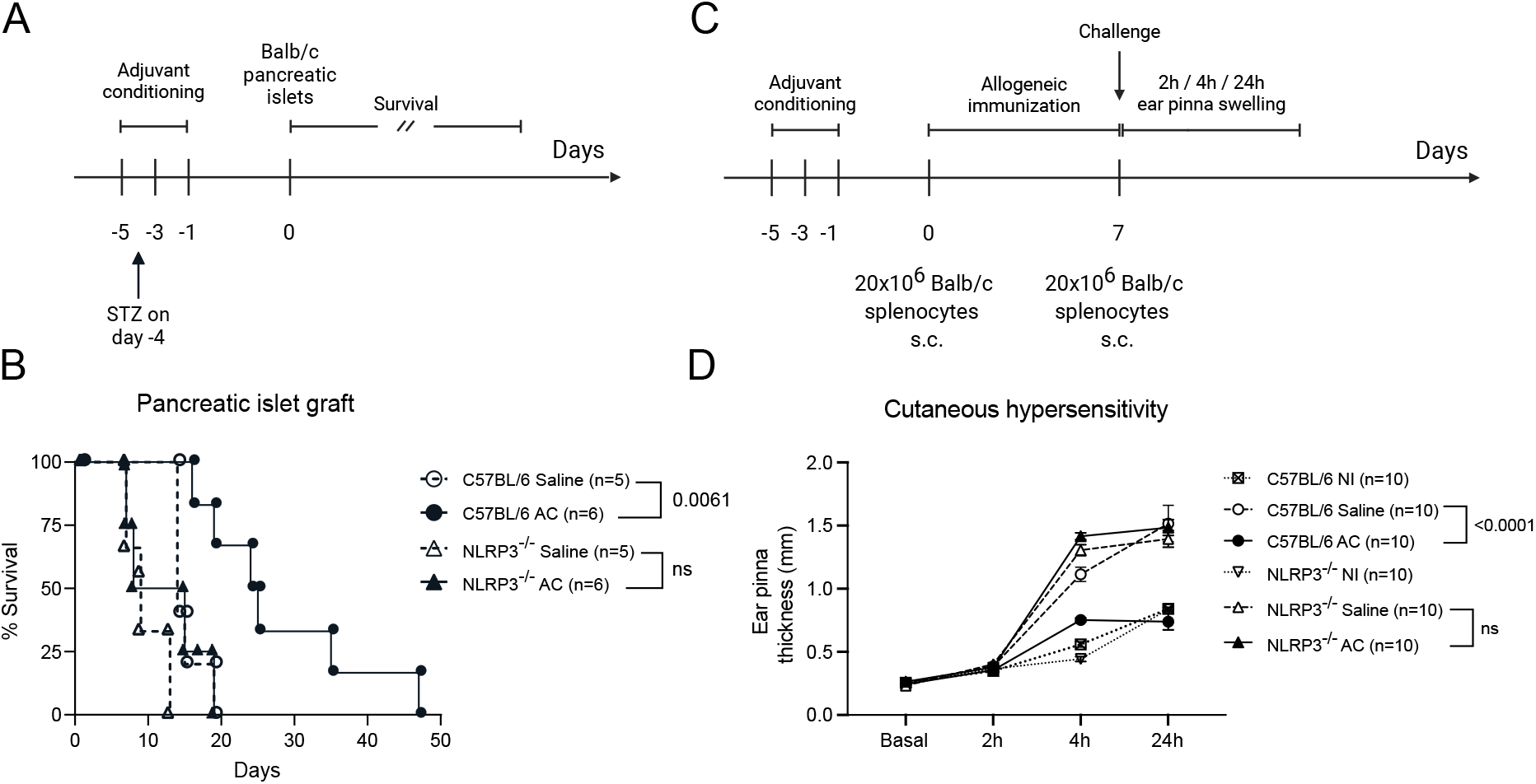
Adjuvant conditioning promotes allogeneic tolerance dependent on NLRP3. **A:** Allogeneic pancreatic islet transplantation experimental design. Briefly, alum or saline-treated C57BL/6 or NLRP3^-/-^ mice were treated with streptozotocin (STZ) 4 days before the transplantation with Balb/c isolated pancreatic islets. Survival of the graft was assessed daily by checking blood glucose levels. **B:** Survival curve of alum or saline-treated C57BL/6 or NLRP3^-/-^ mice that underwent allogeneic pancreatic islet transplantation. C57BL/6 saline (n=5), AC (n=6), NLRP3^-/-^ saline (n=5), AC (n=6). Graft survival significance was assessed by Kaplan-Meier/Mantel-Cox log-rank test. **C:** Cutaneous hypersensitivity experimental design. Briefly, alum or saline-treated C57BL/6 or NLRP3^-/-^ mice were subcutaneously immunized with 20× 10^6^ Balb/c splenocytes on the neck. Seven days later, mice were challenged with 20×10^6^ Balb/c splenocytes injected subcutaneously into the base of the ear, as described by Zecher et al. (2009) (Zecher et al., 2009). **D:** Cutaneous allogeneic response was assessed by measuring ear pinna swelling (mm) at 2-, 4- and 24- hours post-challenge. P-values show the comparison between ear swelling at 24-hours post-challenge. n=10 for each group. Data shown represent three or more experiments and are expressed as mean±SEM; Student t-test was used for analysis; P values are indicated in each graph. All statistical analyses were performed using GraphPad Prism™.

We further investigated the impact of adjuvant conditioning on the allogeneic response using an additional model. C57BL/6 mice were subcutaneously immunized with Balb/c splenocytes, followed by a challenge with Balb/c splenocytes injected into the base of the ear seven days later (Fig 7C), as done before (Zecher et al., 2009). The cutaneous allogeneic response was assessed by measuring ear pinna swelling up to 24 hours post-challenge. Conditioned mice exhibited reduced ear swelling compared to saline-treated controls (Fig 7D), confirming the suppressive effect of the conditioning protocol on the allogeneic response. Consistent with our allogeneic islet transplantation experiments, conditioned NLRP3-/-mice failed to suppress the allogeneic response, showing similar levels of ear swelling as saline-treated controls (Fig 7D). Collectively, the data confirm the critical role of NLRP3 in regulating allogeneic responses in vivo following AC.

### Alum stimulation induces trained immunosuppression in human PBMCs in vitro

Finally, we investigate whether adjuvant conditioning-induced trained immunosuppression also occurs in humans, we adapted an innate training protocol for in vitro studies (Röring et al., 2024; Vuscan et al., 2024). Human PBMCs were stimulated with alum (once, twice, or three times before LPS stimulation (Fig 8A). The supernatant was collected for viability assays and cytokine detection. We observed that a single alum stimulation induced an increase in nearly all the inflammatory cytokines tested after LPS stimulation. When cells were treated with multiple doses of alu, resembling AC, the secretion of the proinflammatory cytokines, and especially IL-12, TNF-αα and IL-6, declined (Fig 8C), and this effect depended on the dose of alum (Fig 8D). Viability assays confirmed cell survival 24 hours after each alum stimulation (Fig 8B). Notably, the same pattern was observed for AC-induced murine MDSCs (Fig 5B-E and 5K-L). These findings demonstrate that AC drives trained immunosuppression both in mouse and human cells.

**Figure 8:**
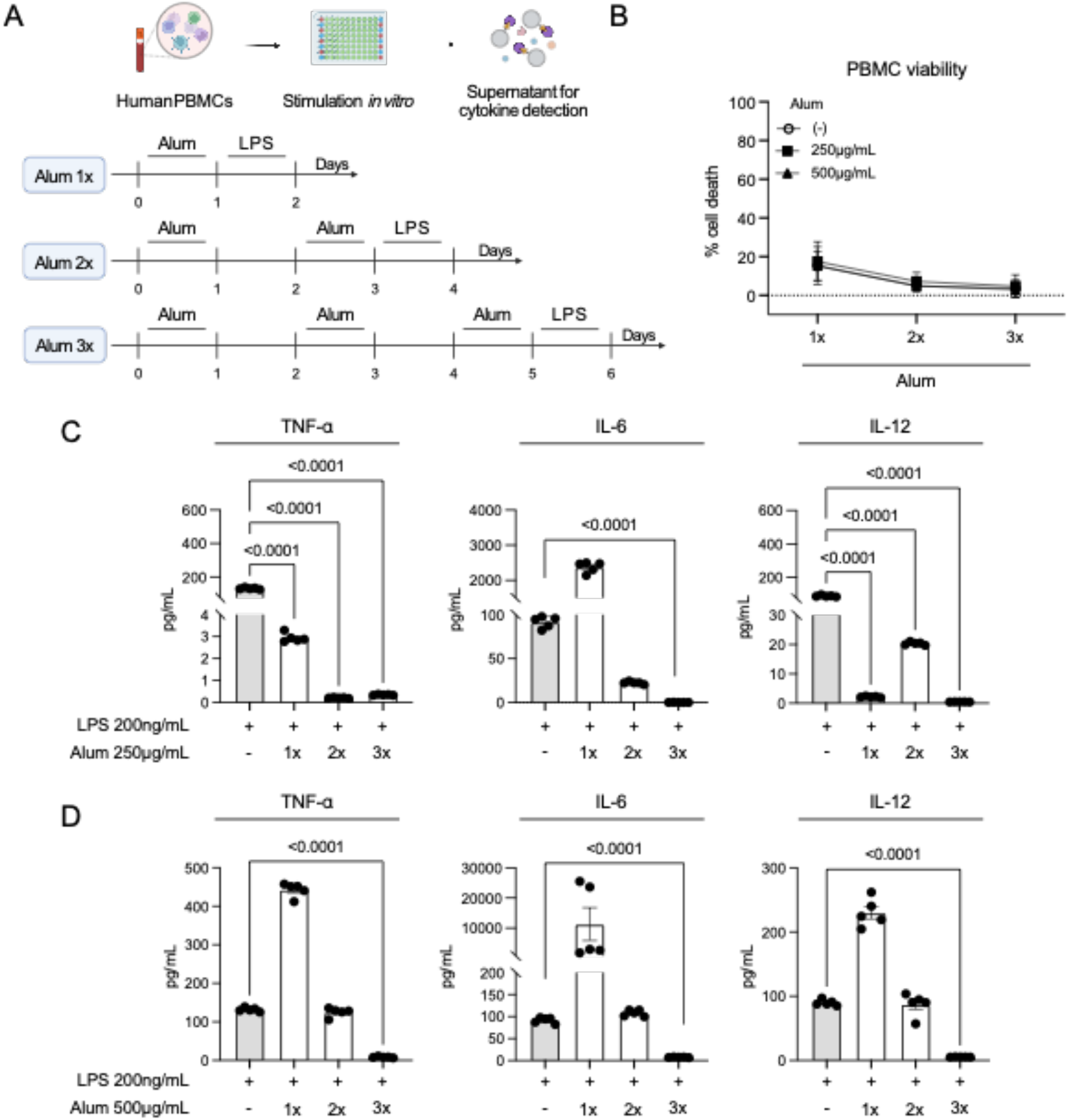
Alum stimulation induces trained immunosuppression in human PBMCs *in vitro*. **A:** Experimental design. Briefly, 3×10^5^ total PBMCs from five different healthy donors were stimulated with alum (250μg/mL and 500μg/mL) once, twice or three times before stimulation with LPS (200ng/mL) for 24h. At the designated times, the supernatant was collected for cytokine measurement using LEGENDplex™ Human Inflammation Panel 1. **B:** Viability of PBMCs stimulated with alum (250μg/mL and 500μg/mL) once, twice or three times before stimulation with LPS (200ng/mL), measured by LDH release. **C:** TNF-α, IL-6 and IL-12 production by PBMCs stimulated with alum (250μg/mL) once, twice or three times before stimulation with LPS (200ng/mL). **D:** TNF-α, IL-6 and IL-12 production by PBMCs stimulated with alum (500μg/mL) once, twice or three times before stimulation with LPS (200ng/mL). Figure created with BioRender.

## Discussion

Our study reveals that adjuvant conditioning expands the population of MDSCs which inhibit antigen-specific T cell proliferation in both in vitro and in vivo settings. Furthermore, we have demonstrated a crucial role for NLRP3 activation and IL-1 signaling in facilitating the expansion and suppressive function of these MDSCs, thereby uncovering a previously unrecognized immunosuppressive pathway triggered by inflammasome activation, possibly related to the process known as trained immunity (Netea et al., 2020). Significantly, in the absence of NLRP3 and IL-1 signaling, the immunosuppressive impact of adjuvant conditioning diminishes, suggesting a potential role for inflammasome activation in fostering allogeneic tolerance in vivo and for the suppressive trained immunity reprograming of MDSCs (Netea et al., 2020; Röring et al., 2024). Furthermore, the NLRP3/IL-1 pathway leads to trained immunosuppressive responses or trained tolerance in vivo. Human PBMCs treated with alum in vitro, show reduced capacity to secrete inflammatory cytokines, suggesting that trained immunosuppression may also be observed in humans. These findings suggest a role for further characterization of immunomodulatory adjuvants that can be used to create a suppressive milieu within a transplant patient.

Currently, several strategies are employed to induce immunosuppression in patients with autoimmune diseases or undergoing transplantation, including various clinical trials testing new drugs and cell-based therapy protocols (McInnes and Gravallese, 2021; Pilch et al., 2021; Ott and Cuenca, 2023). Exploring the potential of FDA-approved adjuvants such as alum represents a novel approach to these protocols. These substances, traditionally known for enhancing immune responses to foreign antigens, could also be leveraged to expand immunosuppressive cell populations, potentially aiding in the induction of tolerance. Our group has previously demonstrated the potential of aluminum hydroxide salts in the protection of mice from neonatal sepsis, inducing emergency myelopoiesis (Rincon et al., 2018), and prolonging allogeneic graft survival (Ge et al., 2023). Here, we demonstrated that adjuvant conditioning expands functional MDSCs which effectively suppress antigen-specific adaptive response both in vitro and in vivo supporting the concept that AC induces trained immunosuppression, or trained tolerance.

The effects of adjuvant conditioning before immunization and/or infections have been studied by other groups, and the mechanism used by each adjuvant is different. In a protocol similar to what we describe here as adjuvant conditioning, EPS, an exopolysaccharide derived from Bacillus subtilis, has demonstrated protective effects in various contexts: preventing allergic eosinophilia (Swartzendruber et al., 2019; Jones and Knight, 2012; Jones et al., 2014), combating infections caused by C. rodentium (Paynich et al., 2017) and S. aureus (Paik et al., 2019), and mitigating severe GVHD (Kalinina et al., 2021). EPS exerts its immunosuppressive actions through a mechanism involving TLR4-dependent IDO expression in dendritic cells (Zamora-Pineda et al., 2023; Kalinina et al., 2024). Similarly, CpG, another TLR ligand, when administered during conditioning prior to immunization, reduces total IgG titers (Wingender et al., 2006; Schmoeckel et al., 2018) and suppresses CD8 T cell cytotoxic responses via an IDO-dependent mechanism (Wingender et al., 2006), highlighting a consistent pattern of TLR-induced immunosuppressive mechanisms. Furthermore, aluminum hydroxide salts have been noted for their ability to suppress antibody production, either alone or in combination with CpG (Schmoeckel et al., 2018).

Previous studies have shown consistent findings regarding inflammasome activation occurring prior to immunization or during infections. Inflammasomes are multi-protein complexes that assemble upon cytosolic detection of microbial patterns or harmful stimuli, resulting in activation of caspase-1, pyroptotic cell death, and the release of IL-1β and IL-18. These processes play a critical role in the control of bacterial and parasitic infections (Lamkanfi and Dixit, 2014; Rathinam and Fitzgerald, 2016; Broz and Dixit, 2016). Despite their association with increased inflammation, inflammasome activation has been reported to exert anti-inflammatory effects as well. Notably, NLRP3 activation has been linked to immunosuppressive outcomes such as reduced CD8 T cell responses against tumors (Daley et al., 2017; Das et al., 2020; Theivanthiran et al., 2020; van Deventer et al., 2023) and induction of protective responses in colitis (Yao et al., 2017; Mak’Anyengo et al., 2018). Activation of caspase-1, a key effector of the inflammasome complex, has been observed to contribute to CD4 T cell depletion during HIV infection (Monroe et al., 2014), while blocking IL-18 signaling in T cells may exacerbate disease severity in murine colitis models (Holmkvist et al., 2016; Mak’Anyengo et al., 2018). Our findings, showing reduced expansion of specific CD4 T cells and diminished antibody production following the conditioning regimen, are consistent with these documented effects in the literature. NLRP3-dependent IL-1 has been implicated in the generation, expansion, and suppressor function of MDSCs in mouse tumor models (Elkabets et al., 2010; Tu et al., 2011; Kaplanov et al., 2019; Koehn et al., 2019; Tengesdal et al., 2021; Papafragkos et al., 2022; Shi et al., 2022; van Deventer et al., 2023). Our study underscores the critical role of NLRP3 activation and IL-1 signaling in promoting both the expansion and suppressive function of these MDSCs in vitro and in vivo, revealing a novel immunosuppressive pathway associated with inflammasome activation, akin to what we term ‘trained immunosuppression or tolerance’. The effects of AC in shaping the adaptive immunity may also be related to a type of antigen-specific training of the innate system, since it leads to long term effects in both immunization protocols and allograft rejection. Inflammasomes have been linked to processes resembling trained immunity, primarily mediated through IL-1 signaling pathways impacting immunometabolism, such as glycolysis (Kol et al., 1997; Tan et al., 2018; Riera et al., 2007), oxidative metabolism (Zhou et al., 2020; Batista et al., 2021), and epigenetic modifications involving histones and DNA (Li et al., 2020; Bellavia et al., 2022), as well as hematopoietic stem cell expansion (Mitroulis et al.).

Moreover, the absence of NLRP3 results in the loss of immunosuppressive reprogramming in both monocyte-MDSCs and PMN-MDSCs in mouse tumor models, underscoring the indispensable role of this well-established inflammatory pathway in immune suppression (Papafragkos et al., 2022). TLR ligands have also been linked to trained immunity processes. It has been observed that mice conditioned with CpG show an expansion of a distinct type of neutrophils that produce fewer extracellular traps, resulting in reduced organ damage in sepsis models (Ng et al., 2020, 2023). This suggests that these neutrophils may undergo a similar reprogramming process as the MDSCs demonstrated in our study. In vitro stimulation of human PBMCs with alum show reduced inflammatory cytokine secretion, indicating a similar mechanism of trained immunosuppression in human cells.

Overall, our findings show how an adjuvant used mainly for boosting the immune response, can also drive immunosuppressive pathways, leading to delayed allograft rejection, while shaping the adaptive immune response. These findings advocate for developing immunomodulatory adjuvants that utilize the NLRP3/IL-1 pathway to induce a suppressive environment in transplant recipients.

## Materials and Methods

### Mice

For the study, 8–10-week-old males and females C57BL/6J (H-2b), Balb/C (H-2d), OT-II (B6.Cg-Tg(TcraTcrb)425Cbn/J), and NLRP3-/-(B6.129S6-Nlrp3tm1Bhk/J) mice were purchased from Jackson Laboratory. All mice were kept under specific pathogen free facility at Boston Children’s Hospital. All mouse experimental protocols were approved by the Institutional Animal Care and Use Committees of Boston Children’s Hospital.

### Drug administration

For adjuvant administration, mice received either saline or alum (8mg in 200 □L) (Thermo Fischer Scientific) intraperitoneally every other day for three times. For IL-1 signaling blockade, KINERET® (Anakinra) was injected intraperitoneally at a dose of 30mg/kg at time points described in each figure.

### MDSCs enrichment

CD11b+GR1+ cells were isolated using negative selection from mouse spleens using the EasySep mouse MDSC (CD11b+Gr1+) isolation kit (StemCell) according to the manufacturer’s protocol. Purity of isolated MDSCs were verified using flow cytometry to be >90%. Isolated MDSCs were cultured in RPMI 1640 medium (ThermoFisher Scientific) containing 10% fetal bovine serum (Corning), 200μg/mL penicillin (ThermoFisher Scientific), 200 U/mL streptomycin (ThermoFisher Scientific), and 0.05mM 2-mecaptoethanol (Sigma-Aldrich) for 24h in the presence or not of LPS (200ng/mL) (Sigma-Aldrich cat 297-473-0).

### CD4 T cell suppression assay

To assess MDSCs suppressor function, naïve CD4+ T cells were isolated from mouse spleens using EasySep mouse naïve CD4+ T cell isolation kit (StemCell) and labeled with CellTrace Violet (CTV; ThermoFisher Scientific). CD11b+GR1+ cells were isolated from alum-treated or saline-treated animals and cultured with naïve CD4 T cells in the presence of 1μg/mL of OVAp 323-339 (InvivoGen) and C57BL/6 total splenocytes for 72h.

### OVA immunization

Wild-type C57BL/6J or NLRP3 deficient mice were adoptively transferred with 106 total splenocytes from OT-II mice intravenously. 24h after the adoptive transfer, mice were immunized subcutaneously with OVA grade-V (100mg) (Sigma-Aldrich) adsorbed either to Alum Imject (1.5mg), or to Resiquimod (R848) (InvivoGen) (50μg) in a homologous prime and boost (day 0 and 7) protocol adapted from Cho et al (2017) (Yi-Li Cho, Michael Flossdorf, Lorenz Kretschmer, Thomas Höfer, Dirk H. Busch, Veit R. Buchholz 2017). At day 15 after priming, mice were euthanized, and spleens were collected for phenotyping and restimulation in vitro with 2μg/mL of OVAp 323-339, and blood was collected for serum OVA-antibody titration. For MDSCs adoptive transfer experiments, 2×106 of purified CD11+GR1+ were injected intravenously 24h before the OVA immunization protocol.

### Flow cytometry and protein expression

Single-cell suspensions were prepared from mouse splenocytes after RBC lysis using ACK lysis buffer (ThermoFisher Scientific). The following antibodies were used for surface staining: CD11b (clone: M1/70; Invitrogen), Ly6C (clone: HK1.4; BioLegend), Ly6G (clone: 1A8; BioLegend), CD3 (clone: 17A2; Invitrogen), CD4 (clone: GK1.5; Invitrogen), program cell death protein ligand 1 (PD-L1; clone: 10F.9G2; BioLegend). Samples were fixed and permeabilized with Fix/Perm buffer according to the manufacturer’s instruction (eBioscience) before intracellular protein staining. The following antibodies were used for intracellular staining: Tbet (clone: 4B10; BioLegend), GATA3 (clone: 16E10A23; BioLegend), RORgT (clone: Q31-378; BioLegend), Foxp3 (clone: MF-14; BioLegend), IFN-g (clone: AN-18; BioLegend), IL-4 (clone: 11B11; BioLegend), IL-17A (clone: TC11-18H10.1; BioLegend), LAP (clone: TW7-16B4; BioLegend). Splenocytes were cultured with GolgiPlug (BD Biosciences) for 48h for intracellular cytokine detection. Cell viability was determined Fixable Viability Dye eFluor™ 780 (eBioscience™), Zombie Red™ or Zombie Aqua™ Fixable Viability (Biolegend). Samples were collected on LSRFortessa HTS with BD FACSDiva v8.0.2 software (BD Biosciences). FlowJo v10 was used for flow data analysis. IDO expression was determined using Purified anti-IDO-1 (clone: 2E2/IDO1; BioLegend).

### Human PBMC isolation for stimulation in vitro

PBMCs from healthy donors were isolated using Lymphoprep density gradient medium centrifugation (Stemcell Technologies). Blood was diluted in a 1:1 ratio with PBS 1x + 2% FBS + 4 mM EDTA to prevent cell death and clumping. The diluted blood was carefully layered on 10 ml of Lymphoprep medium at room temperature in a 50 mL Falcon tube, avoiding mixing. The tube was then placed in a centrifuge and spun for 30 minutes at 900xg at 23 °C with minimum deceleration. After centrifugation, the cell ring between the upper layer (plasma) and the lower layer (Lymphoprep) was collected and washed with PBS 1x + 2% FBS + 4 mM EDTA. Red blood cells were lysed using 10 ml of ACK lysing buffer (Gibco) for 10 minutes at room temperature. The PBMCs were washed again, resuspended in 5 ml of PBS + 2% FBS + 4 mM EDTA, counted, and kept on ice for further use. Isolated PBMCs were plated 3×105 in each well and stimulated with Alum Imject™ at 250 or 5007g/mL for 24h for one, two or three times, and later stimulated with LPS (200ng/mL) for an additional 24h. In between alum stimulations the supernatant was changed, stored at -80oC, and fresh complete RPMI was added. Cell viability after alum stimulation was assessed by LDH release, measured using CyQUANT™ LDH Cytotoxicity Assay (ThermoFisher Scientific).

### Cytokine detection

Cytokine release in the supernatant was assessed using LEGENDplex™ Mouse Inflammation Panel 1 (740150; BioLegend) and LEGENDplex™ Human Inflammation Panel 1 Standard (740811; BioLegend). Serum cytokines were assessed by Mouse IL-1 beta/IL-1F2 DuoSet ELISA (DY40105; R&D Systems™) and Mouse IL-1 alpha/IL-1F1 Quantikine ELISA Kit (MLA00; R&D Systems™).

### Allogeneic pancreatic islet transplantation

To perform the islet transplant model, recipient mice (C57BL/6J) were injected with streptozotocin (STZ) (Sigma-Aldrich) 240 mg/kg intraperitoneally to induce diabetes. Diabetes induction was confirmed in each mouse if blood glucose level was ≥200 mg/dL for 3 consecutive days. Islets from donor mice (Balb/c) were isolated via pancreas collagenase digestion and density gradient centrifugation as described previously (Schuetz et al. 2017). 250 to 300 islets were transplanted under the renal capsule of recipient mice. Blood glucose level was monitored daily, and rejection was defined when blood glucose ≥200 mg/dL for 3 consecutive days.

### Allogeneic cutaneous hypersensitivity

C57BL/6 mice were immunized with 20 × 106 Balb/c splenocytes subcutaneously on the neck. Seven days later, mice were challenged with 20 × 106 Balb/c splenocytes injected subcutaneously into the base of the ear, as described by Zecher et al. (2009) (Zecher et al. 2009). Allogeneic response was assessed by measuring ear pinna swelling (mm) at 2-, 4- and 24-hours post-challenge.

## Data Analysis and Statistics

Statistical analysis was performed using GraphPad Prism version 10. Two-tailed unpaired Student t test was used to calculate differences between experimental animals. One-way analysis of variance was used for multiple comparisons. Graft survival significance was assessed by Kaplan-Meier/Mantel-Cox log-rank test. P value of <0.05 was considered statistically significant difference.

## Data availability

Data acquired specifically for this study are available within the article itself and the supplementary materials. Experimental protocols and additional details regarding methods employed in this study will be made available through reasonable request with the corresponding author.

## Acknowledgements

This work was support by the Hardy Hendren Faculty Development Fund at Boston Children’s Hospital and the Junior Translational Investigator Service Award from the Translational Research Program at Boston Children’s Hospital.

## Supplemental material

**Fig. S1.**
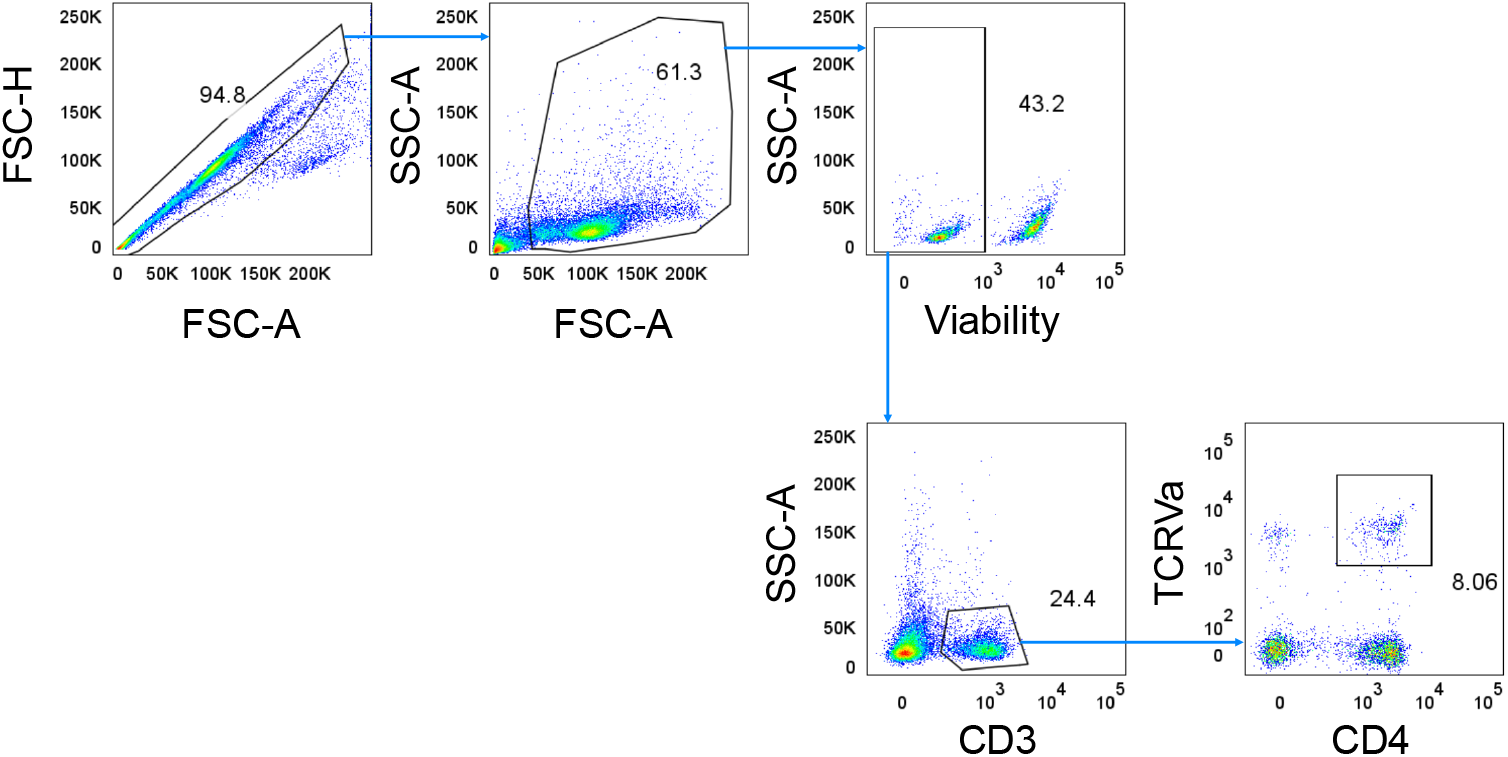
shows the gating strategy used for flow cytometry analysis of immunized C57BL/6 mice in order to find the population of CD4+TCRVα+ cells.

**Fig. S2.**
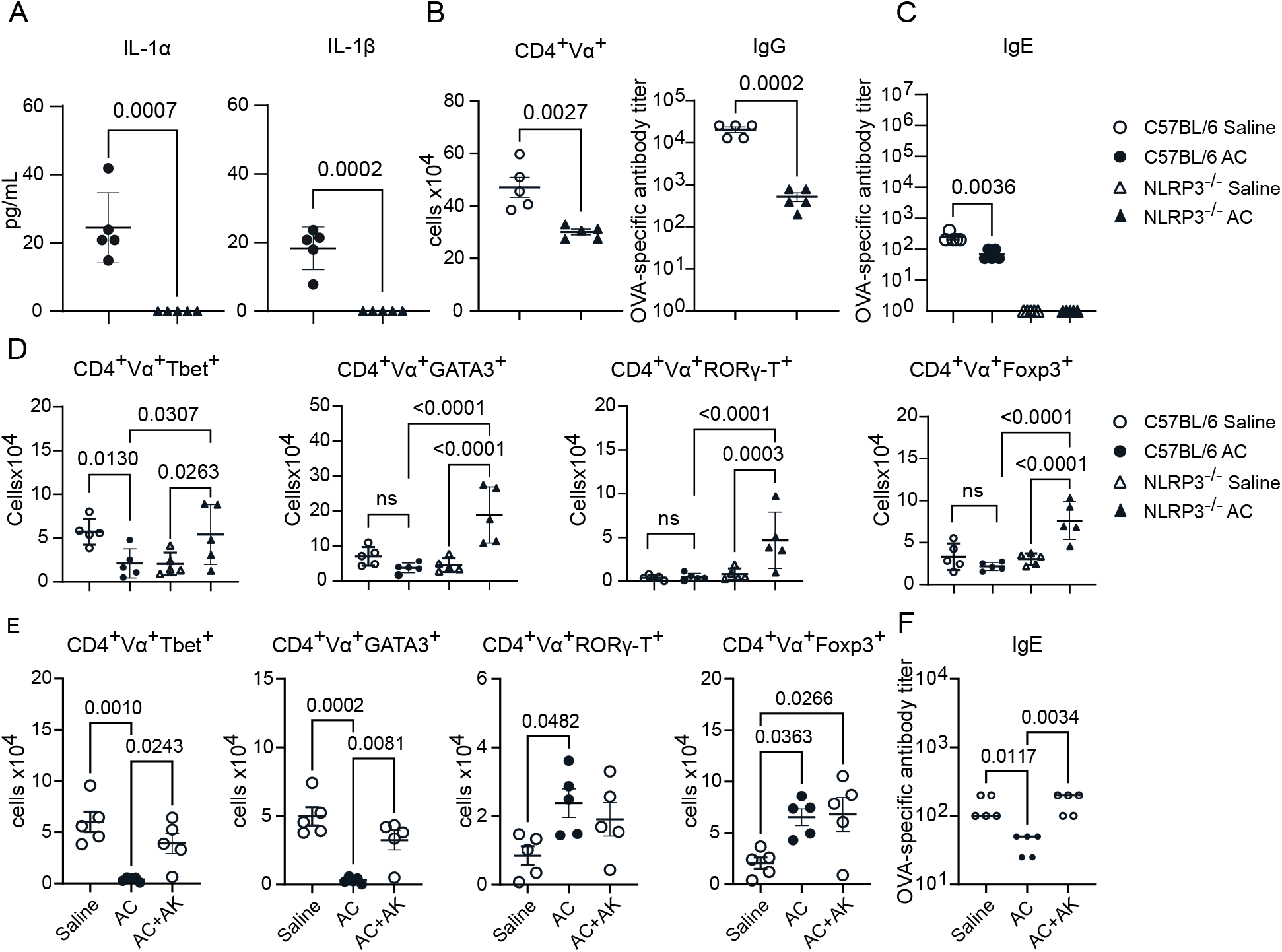
shows serum levels of IL-1α and IL-1β after alum injections; CD4+TCRVα+ cells and IgG levels in C57BL/6 and NLRP3-/-mice after immunization; number of Th1, Th2, Th17, Tregs, and serum IgE levels in C57BL/6 and NLRP3-/-mice after immunization, and mice treated with anakinra prior to immunization.

**Fig. S3.**
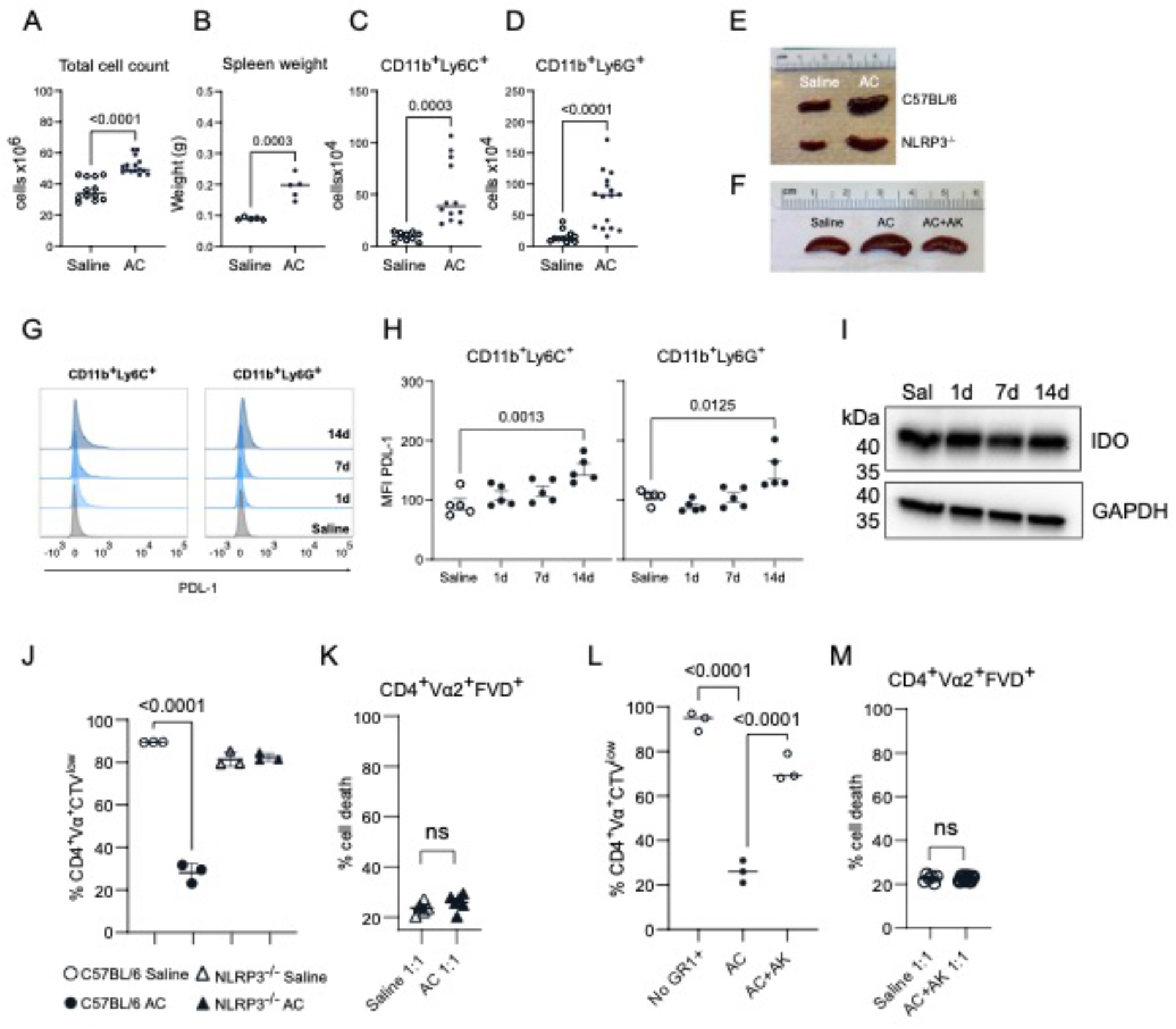
shows total cell count, spleen weight, total MDSC cell count in, PDL-1 and IDO expression in saline or AC-treated C57BL/6 mice; and CD4 T cell proliferation and cell death after culture in vitro with isolated AC-induced CD11b+GR1+ cells from C57BL/6, NLRP3-/- or C57BL/6 mice treated with anakinra during the conditioning protocol.

